# FAM19A5l affects mustard oil-induced peripheral nociception in zebrafish

**DOI:** 10.1101/2020.08.11.245738

**Authors:** Inyoung Jeong, Seongsik Yun, Anu Shahapal, Eun Bee Cho, Sun Wook Hwang, Jae Young Seong, Hae-Chul Park

## Abstract

Family with sequence similarity 19 (chemokine (C-C motif)-like), member A5 (FAM19A5) is a chemokine-like secretory protein recently identified to be involved in the regulation of osteoclast formation, post-injury neointima formation, and depression. Here, we identified *FAM19A5l*, an orthologous zebrafish gene that originated from a common ancestral *FAM19A5* gene. *FAM19A5l* was expressed in trigeminal and dorsal root ganglion neurons as well as distinct neuronal subsets of the central nervous system of zebrafish. Interestingly, *FAM19A5l*^+^ trigeminal neurons were nociceptors that co-localized with TRPA1b and TRPV1, and responded to mustard-oil treatment. Behavioral analysis revealed that the nociceptive response to mustard oil decreased in *FAM19A5l*-knockout zebrafish larvae. In addition, *TRPA1b* and *NGFa* mRNA levels were down- and up-regulated in *FAM19A5l*-knockout and - overexpressing transgenic zebrafish, respectively. Together, our data suggested that FAM19A5l played a role in nociceptive responses to mustard oil by regulating *TRPA1b* and *NGFa* expression in zebrafish.

## INTRODUCTION

FAM19A5/TAFA5 is a member of the family with sequence similarity 19 (chemokine (C-C motif)-like), member A1-5 (FAM19A1-5, also known as TAFA1-5), which encode secretory proteins and are found to be predominantly expressed in the central nervous system (Tom Tang et al., 2004). Previous reports have shown that the FAM19A family is involved in locomotor activity, pain, and fear/anxiety as well as cell migration, survival, proliferation, and differentiation (Wang et al., 2015, Delfini et al., 2013, Kambrun et al., 2018, Shao et al., 2015, Wang et al., 2018a, Choi et al., 2018, Jafari et al., 2019, Lei et al., 2019, Yong et al., 2020, Zheng et al., 2018, Paulsen et al., 2008, Shahapal et al., 2019). Among the members of the FAM19A family, FAM19A5 is known to be highly expressed in several areas of the mammalian brain (Paulsen et al., 2008, Shahapal et al., 2019), suggesting an important role in nervous system development. Recent studies have shown that FAM19A5 inhibits post-injury neointima formation via sphingosine-1-phosphate receptor 2 (Wang et al., 2018b) and osteoclast formation via formyl peptide receptor 2 (Park et al., 2017). A study on FAM19A5-knockout mice has revealed that FAM19A5 is associated with depressive-like and spatial memory-related behaviors (Huang et al., 2020), and a recent report has implicated FAM19A5 in hypothalamic inflammation (Kang et al., 2020). Although FAM19A5 has been associated with nervous system development and psychiatric disorders, its roles in the nervous system remain poorly understood.

Nociception is a sensation induced by noxious stimuli, and numerous studies using several model organisms have shown that transient receptor potential (TRP) ion channels are responsible for detecting a variety of thermal, chemical, and mechanical stimuli (Bandell et al., 2007, Christensen and Corey, 2007, Caterina, 2007, Montell and Caterina, 2007). Zebrafish and other vertebrates perceive noxious stimuli as well as normal sensory stimuli via the trigeminal ganglion (TG) and dorsal root ganglion (DRG) (Bandell et al., 2007, Christensen and Corey, 2007, Caterina, 2007, Montell and Caterina, 2007, Metcalfe et al., 1990, Sagasti et al., 2005, Prober et al., 2008). Zebrafish *Transient Receptor Potential cation channel, subfamily A, member 1 (TRPA1)* and *subfamily V, member 1 (TRPV1)* are known to be expressed in TG neurons and are required for chemo- and thermo-sensation (Prober et al., 2008, Gau et al., 2013). In this study, we identified *FAM19A5l* in zebrafish and demonstrated that its expression in nociceptive trigeminal neurons co-localized with TRPA1b and TRPV1 and responded to mustard-oil treatment. *FAM19A5l* knockout reduced the nociceptive response to mustard oil as well as *TRPA1b* and *NGFa* mRNA expression, whereas its overexpression increased their expression. Therefore, these findings suggest that FAM19A5l plays a role in chemo-sensation by regulating TRPA1b and NGFa expression.

## RESULTS

### Evolutionary history of the *FAM19A* gene family and *FAM19A5l* expression in the zebrafish nervous system

To explore the origin and evolutionary history of the *FAM19A5l* gene, we performed phylogenetic and synteny analyses using the amino acid sequences of *FAM19A* family genes from zebrafish and representative vertebrate species, proposing that the *FAM19A5/5L, FAM19A1*, and *FAM19A2/3/4* lineages may have arisen via local gene duplications in an ancestral gene within *VAC D*. Whole genome duplication (1R and 2R) resulted in *FAM19A5* and *FAM19A5l* being located on *GAC D0* and *GAC D2*, respectively, while the teleost-specific 3R event generated zebrafish *FAM19A5a* and *FAM19A5b* (Supplementary Fig. 1). RT-PCR analysis detected *FAM19A5l* mRNA from development through to the adult brain (Fig. 1A), similar to mammalian *FAM19A5* expression (Tom Tang et al., 2004). Whole-mount *in situ* RNA hybridization revealed that *FAM19A5l* was expressed in various areas of the nervous system, including the subpallium, pallium, dorsal thalamus, preoptic region, midbrain tegmentum, hypothalamus, cerebellar plate, optic tectum, medulla oblongata, spinal cord, retina brain, and spinal cord at 2 to 7 days post-fertilization (dpf; Fig. 1B-H’). The injection of *5xuas:FAM19A5l:mcherry* recombinant DNA into *Tg(huc:gal4vp16);Tg(uas:egfp)* embryos, which express EGFP in post-mitotic neurons (Park et al., 2000), revealed that the majority of *FAM19A5l-mCherry* proteins were localized extracellularly near DAPI^+^/HuC^+^ neurons in the nervous system, indicating that FAM19A5l is a secreted peptide in zebrafish (Fig. 1I-K’”), similar to human FAM19A5 (Wang et al., 2018b).

**Figure 1.**
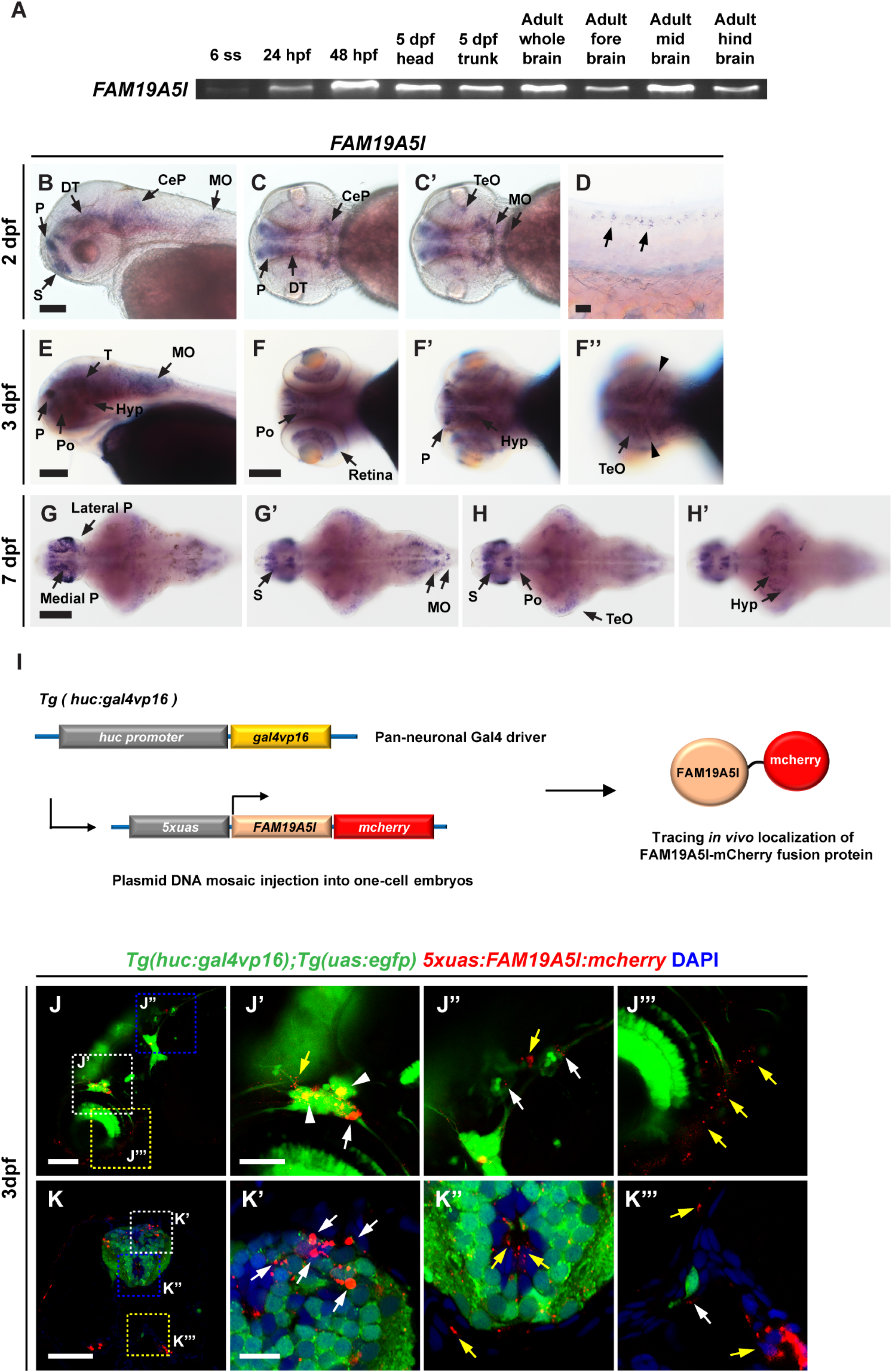
*FAM19A5l* expression in the zebrafish nervous system. (**A**) Reverse transcription-polymerase chain reaction (RT-PCR) of the *FAM19A5l* gene using zebrafish cDNA (ss: somites; hpf: hours post-fertilization; dpf: days post-fertilization). (**B-H’**) *FAM19A5l* RNA expression in larvae and larval brains 2, 3, and 7 dpf: lateral views (B, D, E), dorsal views (C, C’, F-F’’, G, G’) and ventral views (H, H’) of the larvae and brain. Black arrows indicate *FAM19A5l*-expressing areas (CeP-cerebellar plate; DT-dorsal thalamus; Hyp-hypothalamus; MO-medulla oblongata; P-pallium; Po-preoptic region; S-subpallium; T-midbrain tegmentum; TeO-optic tectum). (**I**) Procedure for tracking FAM19A5l proteins. (**J-K’”**): (J-J’’’) Whole-mount *Tg(huc:gal4vp16);Tg(uas:egfp)* 3 dpf larvae injected with *5xuas:FAM19A5l:mcherry*. (K-K’”) Transverse-section views of DAPI-stained 3 dpf *Tg(huc:gal4vp16);Tg(uas:egfp)* larvae injected with *5xuas:FAM19A5l:mcherry*. White arrowheads: GFP^+^/FAM19A5l-mCherry^+^ cells. White arrows: FAM19A5l-mCherry^+^ signals nearby GFP^+^ cell bodies. Yellow arrows: FAM19A5l-mCherry^+^ signals in DAPI^+^/DAPI^-^ in GFP^-^ areas. Scale bar: (B-C’, J’-J’’’, K) 50 μm; (D) 20 μm; (E-H’, J) 100 μm; (K’-K’”) 20 μm.

Using BAC engineering, we generated *Tg(FAM19A5l:egfp-caax)* zebrafish that expressed membrane-bound EGFP protein in *FAM19A5l*-expressing cells (Supplementary Fig. 2). In this transgenic model, *FAM19A5l:*EGFP-CAAX expression was detected in various regions of the brain and spinal cord, similar to endogenous *FAM19A5l* (Supplementary Fig. 2A-Ec). Immunohistochemical (IHC) analysis revealed that *FAM19A5l* was mostly expressed in neurons and radial glial cell subsets in the brain (Supplementary Fig. 2F-Ha’”), whereas in the peripheral nervous system *FAM19A5l*:EGFP-CAAX expression was detected in subsets of TG neurons (Fig. 2A,D, white arrows) and dorsal neurons in the spinal cord (Fig. 2B,E, yellow arrowheads) at 32 hours post-fertilization (hpf) and 3 days post-fertilization (dpf). *FAM19A5l:EGFP-CAAX* was also expressed in retinal ganglion neurons (Fig. 2C, blue arrowheads), subsets of vagal sensory ganglia (Fig. 2D, magenta arrows), and DRG neurons (Fig. 2E, white arrowheads) at 3 dpf. Whole-mount IHC analysis confirmed that *FM19A5l*:EGFP-CAAX^+^ TG and DRG neurons were labeled with Isl1/2^+^ antibodies (Fig. 2F, white arrows, and G, white arrowheads), which are known to mark TG and DRG neurons (Barth et al., 1999, Thisse and Thisse, 2005, Won et al., 2012). Previous reports have demonstrated that *isl2a/2b* is expressed in dorsal longitudinal interneurons (DoLAs) (Tamme et al., 2002) and Rohon-Beard neurons (Appel et al., 1995, Olesnicky et al., 2010) in the spinal cord. Consistently, we observed that *FAM19A5l*:EGFP-CAAX^+^ dorsal neurons were Isl1/2^+^ with small cell bodies and bilateral anteroposterior projections, characteristic of DoLAs, suggesting that these neurons were spinal cord DoLAs (Fig. 2G, yellow arrowheads). Since TG, DRG, and vagal ganglion neurons are sensory neurons (Cho et al., 2002, Huang et al., 2007) and dorsal interneurons are involved in processing sensory inputs in the spinal cord (Jessell, 2000, Lee and Pfaff, 2001), our data suggest that FAM19A5l may play a role in sensation. Using a previously reported FAM19A5 knock-in mouse (Shahapal et al., 2019), we revealed that FAM19A5, the mouse ortholog of zebrafish FAM19A5l, is also expressed in the NF200^+^ DRG (Fig. 2H, H’) and NeuN^+^ dorsal interneurons in the spinal cord (Fig. 2I, I’), suggesting that FAM19A5 expression in sensory neurons is evolutionarily conserved.

**Figure 2.**
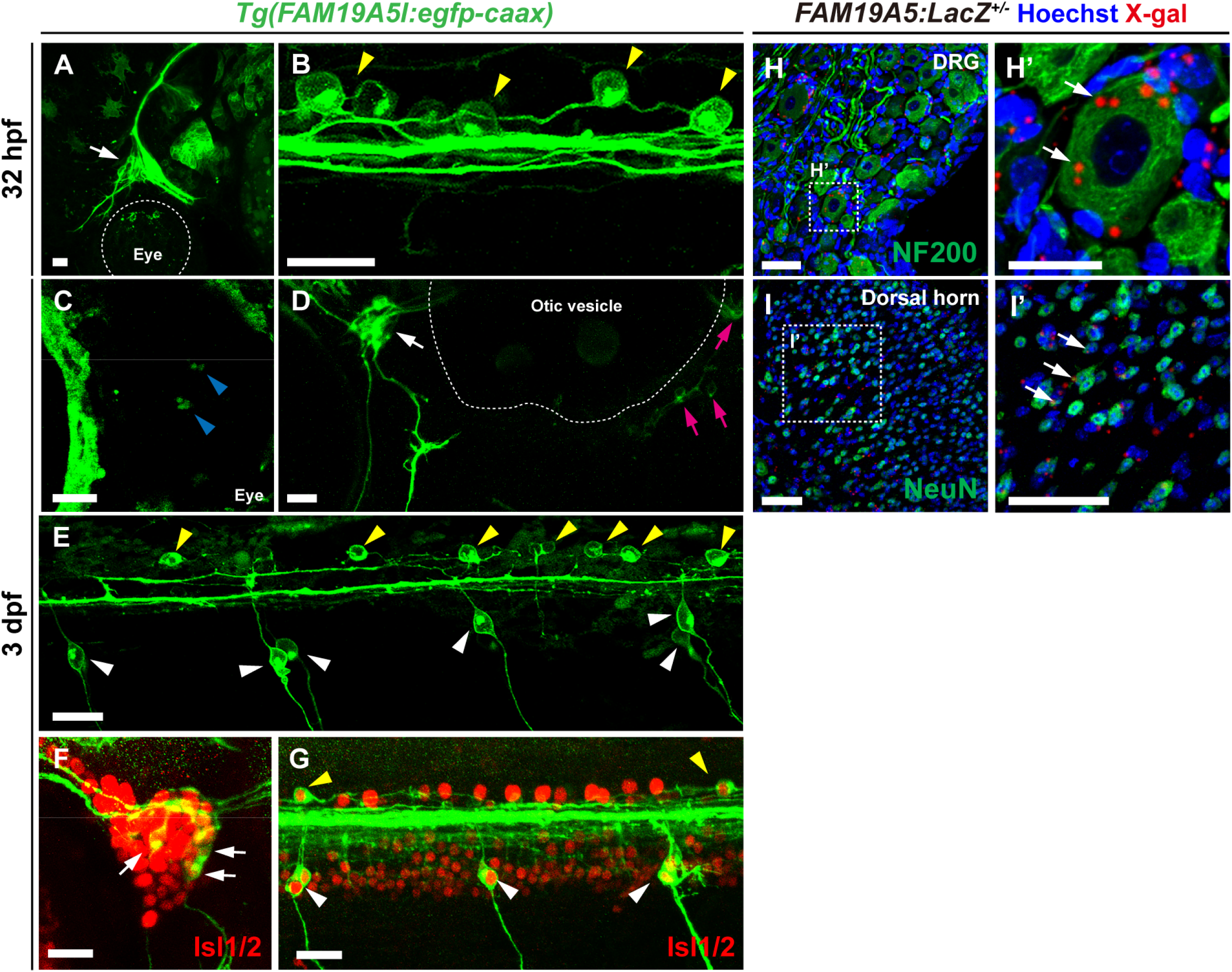
FAM19A5l and FAM19A5 expression in the sensory neurons of zebrafish and mice, respectively. (**A-G**) Lateral view of whole-mount *Tg(FAM19A5l:egfp-caax)* embryos and larvae. White arrows: subsets of trigeminal ganglion (TG) neurons. Yellow arrowheads: subsets of dorsal interneurons. White arrowheads: subsets of dorsal root ganglion (DRG) neurons. Blue arrowheads: subsets of retinal ganglion cells. Red arrowheads: axonal projections. (F-G) TG, DRG, and dorsal interneurons labeled with anti-Isl1/2 antibodies in *Tg(FAM19A5l:egfp-caax)*. White arrows: FAM19A5l^+^/ Isl1/2^+^ TG. White arrowheads: DRG. Yellow arrowheads: dorsal interneurons. Scale bar: 20 μm. (**H-I’**) FAM19A5 expression in adult DRGs and the dorsal spinal cord of FAM19A5:LacZ^+/-^ mice. (H, I) Transverse-section views of DRGs (H) and the dorsal spinal cord (I). IHC using anti-X-gal antibody with neuronal markers; NF200 and NeuN. Scale bar: (A, B, a-a’’, b-b’’) 10 μm; (C, D) 20 μm; (HI’) 50 μm.

### FAM19A5l^+^ TG neurons express *TRPA1b* and *TRPV1* and respond to mustard oil

Since FAM19A5l was expressed in sensory neurons, we hypothesized that FAM19A5l is involved in the response to sensory stimuli. Previous studies have demonstrated that zebrafish *TRPA1* and *TRPV1* are expressed in TG neurons and are required for chemosensation and thermosensation in zebrafish, respectively (Prober et al., 2008, Gau et al., 2013). *In situ* RNA hybridization with *TRPA1b* and *TRPV1* revealed that they were expressed in *FM19A5l*:EGFP-CAAX^+^ TG neurons in *Tg(FAM19A5l:egfp-caax)* zebrafish (Fig. 3A-b’”, white arrows), suggesting a potential role of FAM19A5l in chemosensation and/or thermosensation.

**Figure 3.**
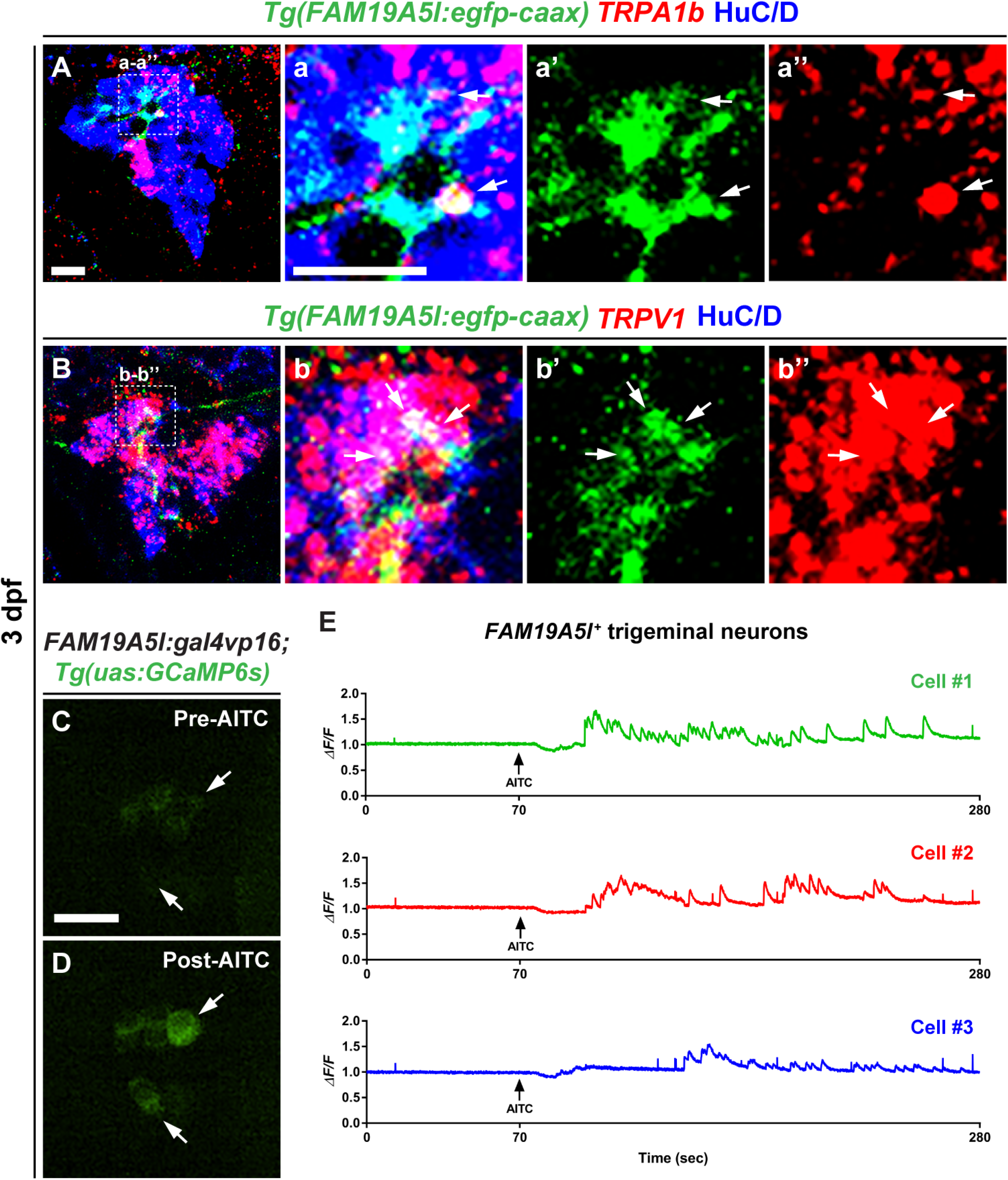
Subsets of FAM19A5l^+^ trigeminal (TG) neurons express *TRPA1b* and *TRPV1* and respond to mustard oil. (**A-b’”**) Lateral view of TGs labeled with *TRPA1b, TRPV1* RNA probes, and anti-HuC/D antibodies in *Tg(FAM19A5l:egfp-caax)*. White arrows: FAM19A5l^+^/HuC/D^+^/*TRPA1b^+^* or *TRPV1^+^* TG. (**C-E**) Calcium imaging using the *FAM19A5l:gal4vp16* BAC DNA micro-injected into *Tg(uas:GCaMP6s)*. AITC: allyl isothiocyanate. White arrows: AITC-responsive FAM19A5l^+^ TG. (E) Three representative calcium imaging traces in FAM19A5l^+^ TG neurons exposed to AITC.

Next, we investigated whether FAM19A5l^+^ TG neurons respond to noxious stimuli using calcium imaging. First, we injected *FAM19A5l:gal4vp16* recombinant BAC DNA into *Tg(uas:GCaMP6s)* embryos to express GCaMP6s, a calcium indicator (Chen et al., 2013), in FAM19A5l^+^ TG neurons. The injected embryos were then treated with allyl isothiocyanate (AITC, mustard oil), a noxious chemical and known TRPA1 agonist (Jordt et al., 2004). Prior to AITC exposure, FAM19A5l^+^ TG neurons rarely showed calcium activity; however, these neurons were activated dramatically after AITC exposure (Fig. 3C-E, Supplementary Video 1). Consistent with previous observations (Esancy et al., 2018), AITC exposure did not elicit synchronized or equivalent calcium responses in FAM19A5l^+^ TG neurons (Fig. 3E). Taken together, these data indicate that FAM19A5l^+^ TG neurons are nociceptors that respond to AITC.

### FAM19A5l is required for AITC-induced nociception but not for heat sensation

To investigate the effect of loss- and gain-of-FAM19A5l function on nociception, we generated *FAM19A5l-knockout* and heat-inducible *Tg(hsp70l:FAM19A5l:p2a-mcherry)* zebrafish (Supplementary Fig. 3). Whole-mount *in situ* RNA hybridization and RT-PCR analysis revealed that *FAM19A5l* expression was decreased and increased in knockout and transgenic zebrafish, respectively (Supplementary Fig. 3). Therefore, we examined the AITC-induced nociceptive response by measuring the locomotor activities of the wildtype, heterozygous, and homozygous siblings of *FAM19A5l*-knockout zebrafish at 3 dpf. Locomotor activity was increased in AITC-treated larvae compared to DMSO-treated larvae (Fig. 4A, B); however, the increase in locomotor activity in *FAM19A5l-*knockout larvae was lower than in wild-type and heterozygous siblings (Fig. 4B), indicating that FAM19A5l function is required for the nociceptive response to AITC. *FAM19A5l*-knockout larvae displayed normal locomotor activity in the absence of AITC, indicating that loss of FAM19A5l function did not affect motor activity (Fig. 4C). Next, we examined whether FAM19A5l overexpression was sufficient to increase the response to the noxious AITC stimulus. However, compared to the DMSO-treated control larvae, AITC treatment increased the locomotor activity of wildtype and heat-shock-induced *Tg(hsp70l:FAM19A5l:p2a-mcherry)* larvae at similar levels (Fig. 4D, E), and the normal locomotor activity of wildtype and transgenic larvae was similar (Fig. 4F). Therefore, these results indicate that FAM19A5l overexpression is not sufficient to augment the nociceptive response to AITC.

**Figure 4.**
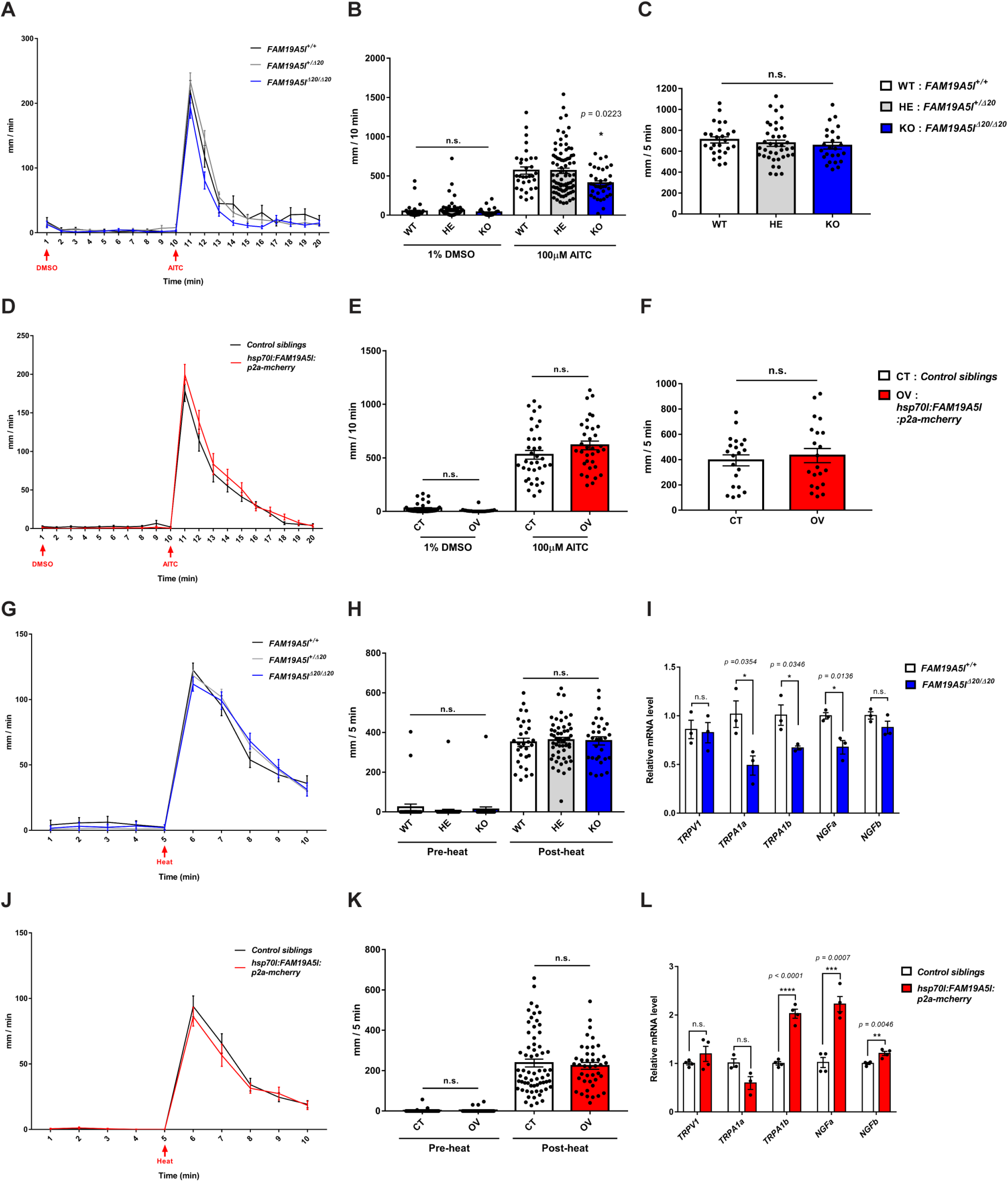
FAM19A5l is required for nociceptive responses to AITC, not heat, and modulates *TRPA1b/NGFa* mRNA expression. (**A,B,D,E,G,H,J,K**) Nociceptive locomotor responses to AITC and heat in *FAM19A5l*-knockout and -overexpressing 3 dpf larvae and their siblings. (A,B,D,E) Distance moved in response to dimethyl sulfoxide (DMSO) and AITC for 10 min. (G,H,J,K) Distance moved in response to heat. (A,D,G,J) Average movement traces for each test. (B,E,H,K) Statistical comparisons between siblings. (A-B) *FAM19A5l*^+/Δ20^ (HE) inbred larvae. *FAM19A5l*^+/+^ (WT; *n* = 31). *FAM19A5l*^+/Δ20^ (HE; *n* = 81). *FAM19A5l*^Δ20/Δ20^ (KO; *n* = 36). Kruskal-Wallis test, Post-hoc Dunn’s multiple comparisons: WT vs KO, \**p* = 0.0470; HE vs KO, \**p* = 0.0404. (D-E) Control siblings (CT; *n* = 35) and *FAM19A5l*-overexpressing larvae (OV; *n* = 34). DMSO: Unpaired *t*-test; AITC: Mann-Whitney *U* test. (G-H) HE inbred larvae. WT (*n* = 30), HE (*n* = 55), and KO (*n* = 30). Oneway ANOVA. (J, K) CT (*n* = 64) and OV (*n* = 42) larvae. Mann-Whitney *U* test. (**C, F**) Distance moved by KO and OV larvae for 5 min at 5 dpf. (C) WT (*n* = 26), HE (*n* = 39), and KO (*n* = 25). One-way ANOVA. (**I,L**) Quantitative RT-PCR in KO and OV 3 dpf larvae and control groups. *β-actin* expression was used to normalize all samples and data represent the results of triplicate/quadruplicate experiments. (I) WT and KO larvae. Unpaired *t*-test. (L) CT and OV larvae. n.s.: not statistically significant.

Since our data showed that FAM19A5l^+^ TG neurons also express TRPV1 (Fig. 3B-B’”), which is required for thermosensation (Gau et al., 2013), we investigated the nociceptive responses to noxious heat in *FAM19A5l-*knockout and -overexpressing transgenic larvae. Although locomotor activity increased in these larvae after heat stimulus over 37 °C at 3 dpf (Fig. 4G,H,J,K), no difference was observed between the increase in knockout and wildtype siblings (Fig. 4G,H,J,K), suggesting that FAM19A5l is not required for the nociception of noxious heat. We confirmed that there was no significant change in the number of TG and DRG neurons in the *FAM19A5l*-knockout or -overexpressing larvae compared to their wildtype siblings (Supplementary Fig. 4A-L). Together, these data suggest that FAM19A5l is involved in nociceptive responses to noxious chemicals, but not heat, in zebrafish.

Interestingly, quantitative RT-PCR (qRT-PCR) revealed that both *TRPA1a* and *TRPA1b* expression decreased by 30–50% in the *FAM19A5l-knockout* larvae compared to the wildtype larvae (Fig. 4I), whereas *TRPA1b* expression was twice as high and *TRPAla* expression did not significantly change in *FAM19A5l*-overexpressing larvae compared to control siblings (Fig. 4L). Consistent with the normal nociception of noxious heat in *FAM19A5l-knockout* and -overexpressing transgenic larvae (Fig. 4H,K), *TRPV1* expression, which is required for thermosensation, did not change in these larvae (Fig. 4I,L).

Since TRPA1 is known to be up-regulated by nerve growth factor (NGF) in rat TG neurons (Diogenes et al., 2007), we hypothesized that FAM19A5l may be involved in the regulation of *NGF* expression. To investigate this possibility, we detected *NGFa* and *NGFb* expression, and found that *NGFa* expression decreased and increased in *FAM19A5l*-knockout and -overexpressing transgenic larvae, respectively, whereas *NGFb* expression increased slightly in the *FAM19A5l*-overexpressing transgenic larvae but was unaffected in *FAM19A5l*-knockout larvae (Fig. 4I,L). Taken together, these data suggest that FAM19A5l is involved in chemosensation by modulating *TRPA1b/NGFa* mRNA expression.

## DISCUSSION

Zebrafish and other vertebrates perceive noxious stimuli as well as normal sensory stimuli such as chemical, thermal, and mechanical stimuli via TG and DRG neurons (Bandell et al., 2007, Christensen and Corey, 2007, Caterina, 2007, Montell and Caterina, 2007, Metcalfe et al., 1990, Sagasti et al., 2005, Prober et al., 2008). TRP ion channels mediate a variety of sensations in response to pH/chemicals, mechanical stimuli, and temperature. Although TRPA1 is known to detect chemical stimuli, its role in the detection of thermal stimuli remains controversial; however, heat-induced TRPV1 responses are conserved in vertebrates (Meents et al., 2019). In this study, we showed that FAM19A5l is expressed in TG neurons and responds to AITC treatment. Behavioral analysis in *FAM19A5l*-knockout zebrafish showed that FAM19A5l is required for the nociceptive response induced by AITC but not by heat, indicating that FAM19A5l plays an essential role in chemosensation. In addition, we revealed that FAM19A5l is required for *TRPA1b* expression but not that of *TRPV1*, consistent with previous reports in zebrafish that TRPA1 is required for chemosensation, but not thermosensation (Prober et al., 2008), whereas TRPV1 is required for thermosensation (Gau et al., 2013).

Previous studies have shown that TRPA1 is expressed in subsets of DRG and TG neurons and that many TRPA1-expressing neurons are peptidergic nociceptors that co-express TRPV1 (Story et al., 2003, Viana, 2016). Moreover, TRPA1 activation in peptidergic nociceptive neurons has been shown to cause the local release of neuropeptides, such as substance P, calcitonin gene-related peptide, or neurokinin A, via Ca^2+^-dependent exocytosis (Gustavsson et al., 2012). Neuropeptide secretion in these nociceptors in turn amplifies nociception by TRPA1 sensitization and neurogenic inflammation mediated by positive feedback regulation (Kadkova et al., 2017). Since FAM19A5l is a secreted neurokine and FAM19A5l^+^ TG neurons co-express TRPA1b and TRPV1, we hypothesized that FAM19A5l may be a neuropeptide released upon TRPA1 activation by noxious stimuli. However, this possibility was ruled out because *TRPA1b* expression was down- and up-regulated at the transcriptional level in FAM19A5l-knockout and -overexpressing transgenic zebrafish, respectively. Thus, FAM19A5l may be an upstream regulator of TRPA1b that is required to induce and maintain TRPA1b expression under normal, unstimulated conditions.

Proinflammatory signals released at the site of injury, such as prostaglandins, bradykinin, and serotonin, are known to act as upstream regulators of TRPA1 activity via G-protein-coupled receptors and phospholipase C-coupled signaling cascades. These mediators can affect the TRPA1 activation threshold to make nociceptors more sensitive to noxious stimulation (Kadkova et al., 2017); however, FAM19A5l appears to modulate TRPA1 expression at the transcriptional level. Consistently, our behavioral study revealed that FAM19A5l-overexpressing transgenic zebrafish displayed normal behavior in the absence of noxious stimuli but increased *TRPA1b* mRNA expression, suggesting that FAM19A5l does not modulate TRPA1 activity.

Interestingly, our data also showed that NGF expression was down- and up-regulated in *FAM19A5l*-knockout and -overexpressing transgenic zebrafish, respectively, suggesting that FAM19A5l is also required to induce and maintain NGF expression under normal, un-stimulated conditions. Although NGF is known to induce TRPA1 expression via receptor tyrosine kinase receptor-mediated neurotrophic signals (Veldhuis et al., 2015, Diogenes et al., 2007), upstream modulators of NGF expression in nociception/pain responses are poorly understood.

In summary, this study identified FAM19A5l in zebrafish and demonstrated that its expression in nociceptive TG neurons co-localized with TRPA1b and TRPV1 and responded to mustard oil treatment. In addition, FAM19A5l knockout reduced the nociceptive response to mustard oil alongside *TRPA1b* and *NGFa* mRNA expression, while its overexpression increased their expression. Therefore, our data suggest that, unlike neuropeptides released from peptidergic nociceptive neurons and proinflammatory signals released at the site of injury, which regulate nociception by modulating TRPA1 activity, FAM19A5l plays a key role in chemosensation by regulating TRPA1b and NGF expression.

## ACKNOWLEDGMENTS

This work was supported by the Collaborative Genome Program of the Korea Institute of Marine Science and Technology Promotion (KIMST) funded by the Ministry of Oceans and Fisheries (MOF; No. 20180430).

## AUTHOR CONTRIBUTIONS

I.J., J.Y.S., and H.C.P. designed the experiments, analyzed data, and wrote the manuscript. I.J., S.Y., A.S., and E.B.C. conducted the experiments. S.W.H. advised and contributed to editing of the manuscript. All authors reviewed the manuscript.

## STAR METHODS

### Resource availability

#### Lead contact and materials availability

Further information and requests for resources and reagents should be directed to and will be fulfilled by the lead contact Hae-Chul Park.

### Experimental model and subject details

All experimental procedures were approved by the Korea University Institutional Animal Care & Use Committee (IACUC; KOREA-2019-0165, KOREA-2016-0091-C3) and performed in accordance with the animal experiment guidelines of Korea National Veterinary Research and Quarantine Service.

For the zebrafish experiments, adult zebrafish and embryos were raised under a 14 h light and 10 h dark cycle at 28.5 °C. Adult zebrafish were fed brine shrimp (*Artemia*; INVE) twice a day. Embryos and larvae were staged by “hour post-fertilization (hpf),” “day postfertilization (dpf)” according to their morphological features (Kimmel et al., 1995), and 6–8 month-old male adult zebrafish were used in the study. At 24 hpf, 0.003 % (w/v) 1-phenyl-2-thiourea (PTU) in embryo medium was used to block pigmentation in zebrafish embryos. Wild-type AB strain, *FAM19A5l^km12^*-knockout, *Tg(FAM19A5l:egfp-caax), Tg(hsp70l:FAM19A5l:p2a-mcherry), Tg(huc:gal4vp16)* (Kimura et al., 2008), *Tg(uas:egfp)* (Asakawa et al., 2008), *nacre (mitfa^b692^* (Lister et al., 1999), received from Koichi Kawakami [NIG, Japan]), and *Tg(uas:GCaMP6s)* (Muto et al., 2017) (received from Koichi Kawakami [NIG, Japan]) zebrafish were used in this study. Transgenic *Tg(FAM19A5l:egfp-caax)* and *Tg(hsp70l:FAM19A5l:p2a-mcherry)* zebrafish were generated in the laboratory by microinjection with linearized *FAM19A5l:egfp-caax* BAC DNA or co-microinjection with *hsp70l:FAM19A5l:p2a-mcherry* plasmid DNA and tol2 transposase mRNA. *FAM19A5l^km12^* wildtype, heterozygotes, and homozygotes were genotyped using PCR using the primers listed in the Key Resources Table.

For experiments involving mouse models, mice were housed under temperature-controlled (22–23 °C) conditions with a 12 h light and 12 h dark cycle (lights on at 8:00 am). The mice were given standard chow and water *ad libitum*. All animal experiments were designed to use the fewest mice possible, and anesthesia was administered. Wildtype C57BL/6J female mice were purchased from Orient Bio (Seongnam, South Korea) and mated with heterozygous *FAM19A5-LacZ KI* males (Shahapal et al., 2019). *FAM19A5l-LacZ* KI mice were genotyped by PCR using the primers listed in the Key Resources Table.

### Method details

#### Data acquisition and phylogenetic analysis

FAM19A family amino acid sequences were retrieved from the human, mouse, anole lizard, chicken, zebrafish, stickleback, tetraodon, and spotted gar genome databases using the Ensembl Genome Browser (http://www.ensembl.org). The amino acid sequences were then aligned using MUSCLE in MEGA 6.06 with the default alignment parameters. A maximum likelihood phylogenetic tree was constructed using the Jones-Taylor-Thornton model in MEGA 6.06 with 100 bootstrapped replicates. Gene orthology or paralogy were further investigated using synteny and search tools in the Ensembl Genome Browser.

#### Synteny analysis and evolutionary history

Synteny analysis was performed by comparing the Contig Views of genome regions containing the peptide and receptor loci. The chromosome localization information of orthologs/paralogs of neighboring genes was obtained from the Ensembl Genome Browser. Chromosome fragments with reliable synteny were matched with the reconstructed vertebrate ancestral chromosome model by Morishita et al. (Nakatani et al., 2007), as described previously (Kim et al., 2014, Yun et al., 2015).

#### Reverse transcription PCR (RT-PCR) and FAM19A5l cloning in zebrafish

To isolate the zebrafish *FAM19A5l* gene, total RNA was extracted from zebrafish embryos, larvae, and adult brains using TRIzol reagent (Thermo Fisher Scientific). Total RNA was synthesized into cDNA using a reverse transcription kit (ImProm-IITM Reverse Transcriptase, Promega), according to the manufacturer’s instructions. RT-PCR was performed for the *FAM19A5l* gene using primer sequences designed using Ensembl (*FAM19A5l*: ENSDARG00000068100). To clone genes, PCR products from 5 dpf cDNA were amplified using PCR and cloned into the pGEM®-T Easy Vector (Promega).

#### Whole-mount RNA in situ hybridization

Cloned vectors were linearized using restriction endonucleases (New England Biolabs) and transcribed using digoxygenin RNA labeling mix (Roche). Whole-mount RNA *in situ* hybridization was performed using two methods: chromogenic reaction with alkaline phosphatase-NBT/BCIP (Roche) (Thisse and Thisse, 2008) and fluorescent reaction with Tyramide Signal Amplification (Perkin Elmer) (Brend and Holley, 2009).

#### Immunohistochemistry in zebrafish

Zebrafish embryos and larvae were anesthetized using 200 mg/L of ethyl 3-aminobenzoate methanesulfonate salt (MS222, Sigma-Aldrich) until movement ceased, fixed in 4 % paraformaldehyde, and embedded in 1.5 % agar blocks containing 5 % sucrose. After equilibration in 30 % sucrose solution, frozen blocks were cut into 14-μm sections using a cryostat microtome and mounted on glass slides. Sections were rinsed several times with phosphate buffered saline (PBS), blocked in 2 % bovine serum albumin with sheep serum, and then treated with primary antibodies overnight at 4 °C. After being washed for 2 h with PBS, the slides were treated with the appropriate secondary antibodies overnight at 4 °C, washed for 2 h with PBS, and mounted. The following primary antibodies were used: mouse anti-HuC/D (1:100, Thermo Fisher Scientific, Cat. No. A21271), mouse anti-Isl1/2 (1:100, Developmental Hybridoma Bank, 39.4D5), rabbit anti-BLBP (1:500, Millipore, ABN14), and chick anti-GFP (1:200, Abcam, Cat. No. AB13970). Alexa 488-, 568-, and 647-conjugated secondary antibodies were used for labeling (1:1000, Thermo Fisher Scientific, Cat. No. A-11001, A-11004, A11008, A-11011, and Abcam Cat. No. ab96947). Nuclei were stained with DAPI (D1306, Thermo Fisher Scientific).

#### X-gal staining and immunohistochemistry in mice

For X-gal staining with multiple fluorescence labeling, 8-week-old male mice were perfused with 4 % PFA in PBS. Their brains were isolated before being post-fixed in the same solution for 3 h, cryo-protected in 30 % sucrose in PBS, and serially sectioned using a cryostat. The spinal cord was cut into 20-μm slices and stored in 50 % glycerol in PBS at −20 °C. For the DRG, 10-μm slices were attached to silane-coated glass slides and stored at −20 °C. For X-gal staining, sections were brought to room temperature, washed three times in PBS for 5 min each, and transferred into X-gal staining solution at 37 °C overnight. After X-gal staining, sections were blocked with 3 % bovine serum albumin and 0.1 % Triton X-100 in PBS for 30 min and incubated with mouse anti-NeuN (1:1000, Millipore, MAB377) and rabbit anti-NF200 (1:1000, Sigma-Aldrich, N4142) primary antibodies overnight at 4 °C.

#### Generation of transgenic zebrafish and plasmid DNA constructs

To generate BAC *Tg(FAM19A5l:egfp-caax)* zebrafish, we used zebrafish bacterial artificial chromosomes (BAC; CH211-240G9) containing the *FAM19A5l* gene. The *FAM19A5l* BAC clone was modified using an *E. coli*-based homologous recombination system to generate *FAM19A5l:egfp-caax* and *FAM19A5l:gal4vp16* BAC DNA. DNAs coding EGFP-CAAX and Gal4VP16 were fused into the ATG site of the *FAM19A5l* open reading frame (ORF) in BAC. Transgenic zebrafish were generated by micro-injecting *FAM19A5l:egfp-caax* BAC DNA into one-cell embryos. To produce *Tg(hsp70l:FAM19A5l:p2a-mcherry)* fish, we amplified the *FAM19A5l* ORF from cDNA using PCR with forward and reverse primers containing attB1 and attB2 sites, as listed in the Key Resources Table. The PCR product containing attB sites was cloned into a middle entry vector using BP clonase (Thermo Fisher Scientific) along with the 5’-entry clone containing the heat shock 70l promoter sequence and the 3’-entry clone containing the mCherry fused viral 2A peptide (Kim et al., 2011). The multi-site gateway LR reaction was performed using LR II clonase with entry clones, according to the manufacturer’s recommendations (Invitrogen). To produce the *5xuas:FAM19A5l:mcherry* plasmid, we performed identical cloning strategies using the 3’-entry clone containing mCherry rather than P2A-mCherry.

#### Generation of FAM19A5l-knockout zebrafish

To generate *FAM19A5l*-knockout zebrafish, we used the CRISPR/Cas9 system, as reported by the Chen et al. (Jao LE, 2013). The Cas9 target site and oligonucleotides for generating the single guide RNA were designed using the ZiFiT web site. The guide RNA vector was cloned and guide RNAs synthesized as described previously (Jao LE, 2013). One-cell stage embryos were injected with a mixture of guide RNA (50 pg) and Cas9 protein (ToolGen, 50 pg) and then raised to adults (F0). To identify founder zebrafish carrying germ-line transmitted mutations, F1 embryos outcrossed from F0 and wild-type zebrafish were collected. Genomic DNA was extracted from 24 hpf F1 embryos, and T7 endonuclease I assays were performed, as described previously (Jao LE, 2013). After candidate founder fish containing mutations were identified, short genomic regions flanking the target site were amplified from the founder genomic DNA using PCR and cloned into the pGEM®-T Easy Vector (Promega). Mutations were analyzed using DNA sequencing. The primers used are listed in the Key Resources Table.

#### Calcium imaging

Briefly, *FAM19A5l:gal4vp16* BAC DNA was micro-injected into one-cell *Tg(uas:GCaMP6s);nacre* embryos. Zebrafish larvae expressing GCaMP6s in TGs were embedded in 2 % low-melting agarose in a dish containing E3 medium with 100 μM D-tubocurarine chloride hydrate (Sigma; T2379). Time-lapse recording was carried out under 50 msec exposure at 20 fps for 280 s using a spinning disc confocal microscope (CSU-X1, Nikon). Larvae were treated with 100 μM allyl isothiocyanate (mustard oil, AITC, 377430, Sigma-Aldrich) at 100 s. The images obtained were analyzed using NIS-Elements software.

#### Larval behavior analyses

Daniovision (Noldus) obtaining automated tracking software Ethovision XT12 (Noldus) was used for all larval behavior analyses. All behavioral experiments were performed from 10:00 a.m.-3:00 p.m. using siblings from a single parent. All larvae were genotyped after experiments to avoid affecting their behavior and biases. Wildtype siblings and *FAM19A5l*-knockout or control and heat-inducible *F4M19A5l*-overexpressing 3 dpf larvae were used for the nociception study. To measure locomotor activity, 5 dpf larvae that were raised under 14/10 h light/dark cycles at 28.5 °C were used. To induce locomotor responses using AITC or a heat stimulus, we exposed larvae to AITC (377430, Sigma-Aldrich) or E3 media over 37 °C, similar to a previously described method (Prober et al., 2008). First, larvae were treated with 1 % dimethyl sulfoxide (DMSO; Biosesang, D1022) followed by 100 μM allyl isothiocyanate in 1 % DMSO after 10 min. Heat-inducible overexpressing larvae were exposed to 39 °C heat shock at 48 hpf for 20 min. All larvae underwent identical testing procedures to assess locomotor activity: 5 dpf larvae were adapted to light conditions for 10 min and then recorded for 5 min under light conditions.

#### Quantitative RT-PCR analyses

Quantitative RT-PCR was performed using Light Cycler 96 Instrument (Roche) with cDNAs synthesized using total RNA extracted from 3 dpf wildtype and *FAM19A5l*-knockout larvae. Each reaction mixture contained 2.5 μL of cDNA as a template, 0.2 μM forward and reverse primers, and 2× Fast Start Essential DNA Green Master Mix (Roche). The following reaction program was used: 95 °C for 10 min followed by 40 cycles of 95 °C for 10 seconds (sec), 60 °C for 10 sec, and 72 °C for 10 sec. We verified gene expression levels of *β-actin*, *FAM19A5l*, *TRPA1a, TRPA1b, TRPV1, NGFa*, and *NGFb*. The primer sequences used are listed in the Key Resources Table.

### Quantification and statistical analysis

All statistical analyses were performed using GraphPad Prism 7 software (GraphPad Software). For comparisons between two groups, Student’s unpaired *t*-test and Mann-Whitney *U* test were used when data were or were not normally distributed, respectively. For comparisons among three groups, one-way analysis of variance (ANOVA) and the Kruskal-Wallis test with post-hoc analysis were performed when data were or were not normally distributed, respectively. Two-tailed *P* values of less than 0.05 were considered statistically significant. Graph bars and error bars represent the mean ± standard error of the mean (SEM).

### Data and code availability

Data supporting the findings of this study are available in the article and its Supplementary Information files, or from the corresponding authors on reasonable request. All analyses were performed using NIS-Elements software, Prism7, MEGA6.06, and Daniovision, as indicated in the Results and Methods sections.

### Key Resources Table

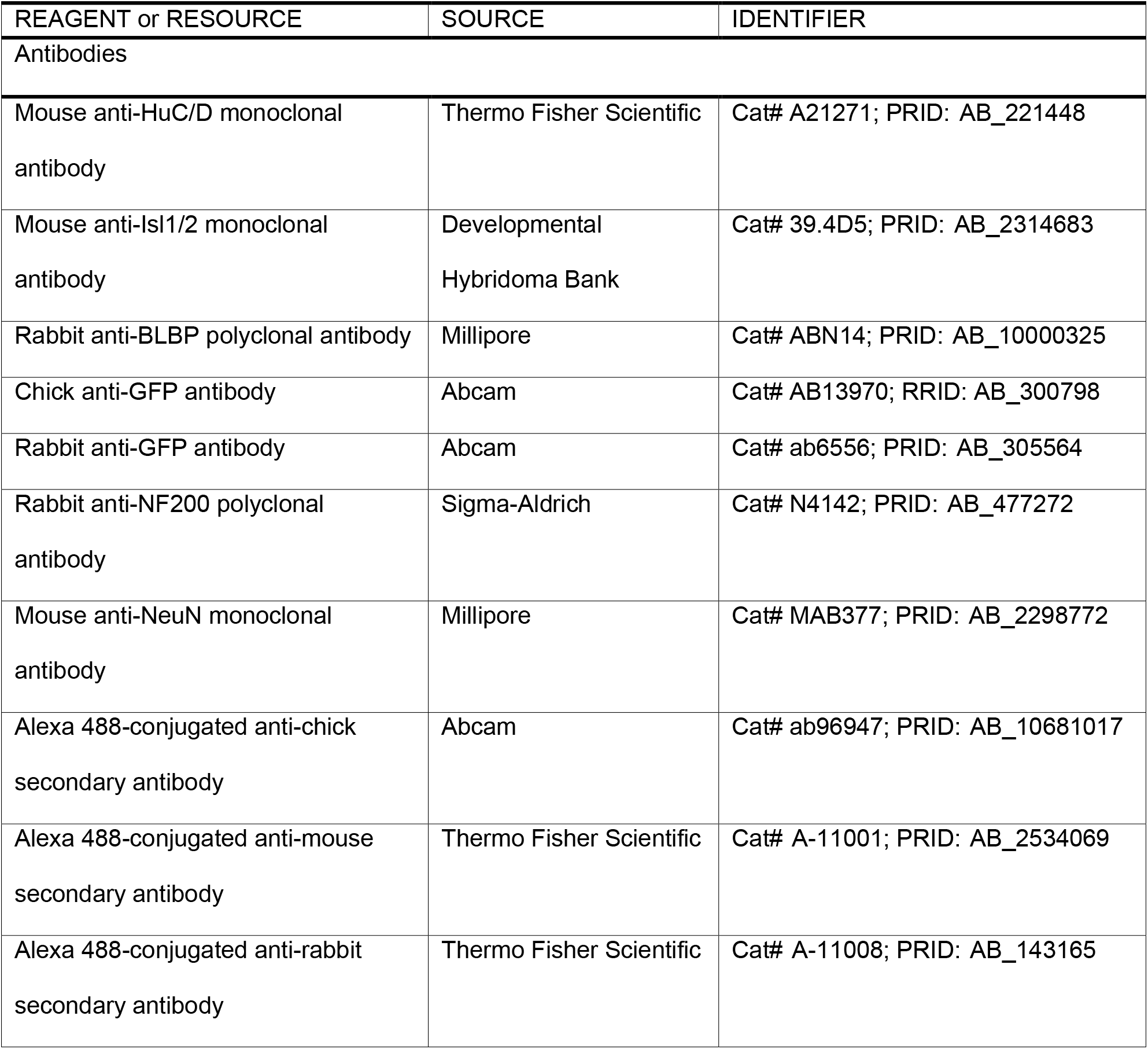
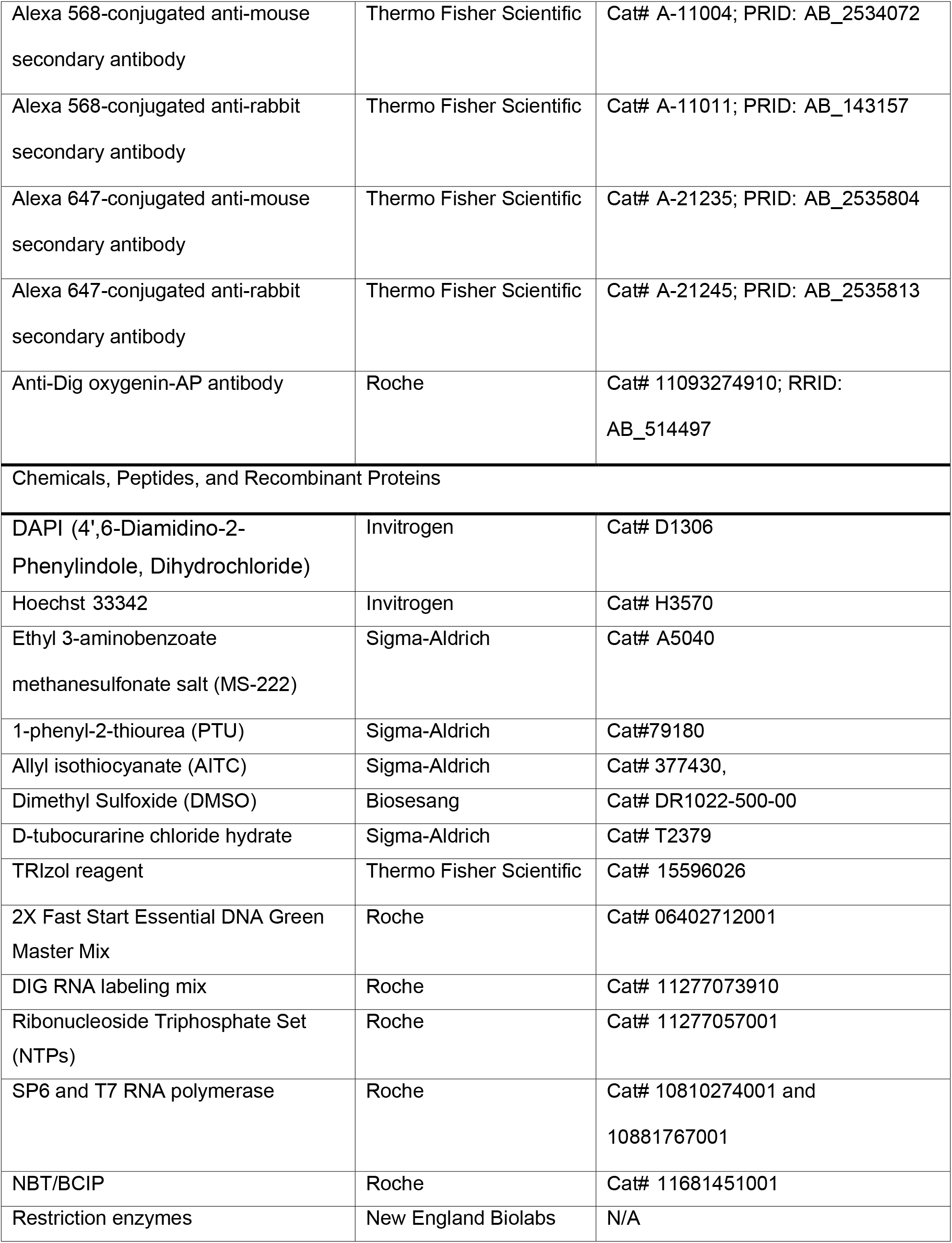
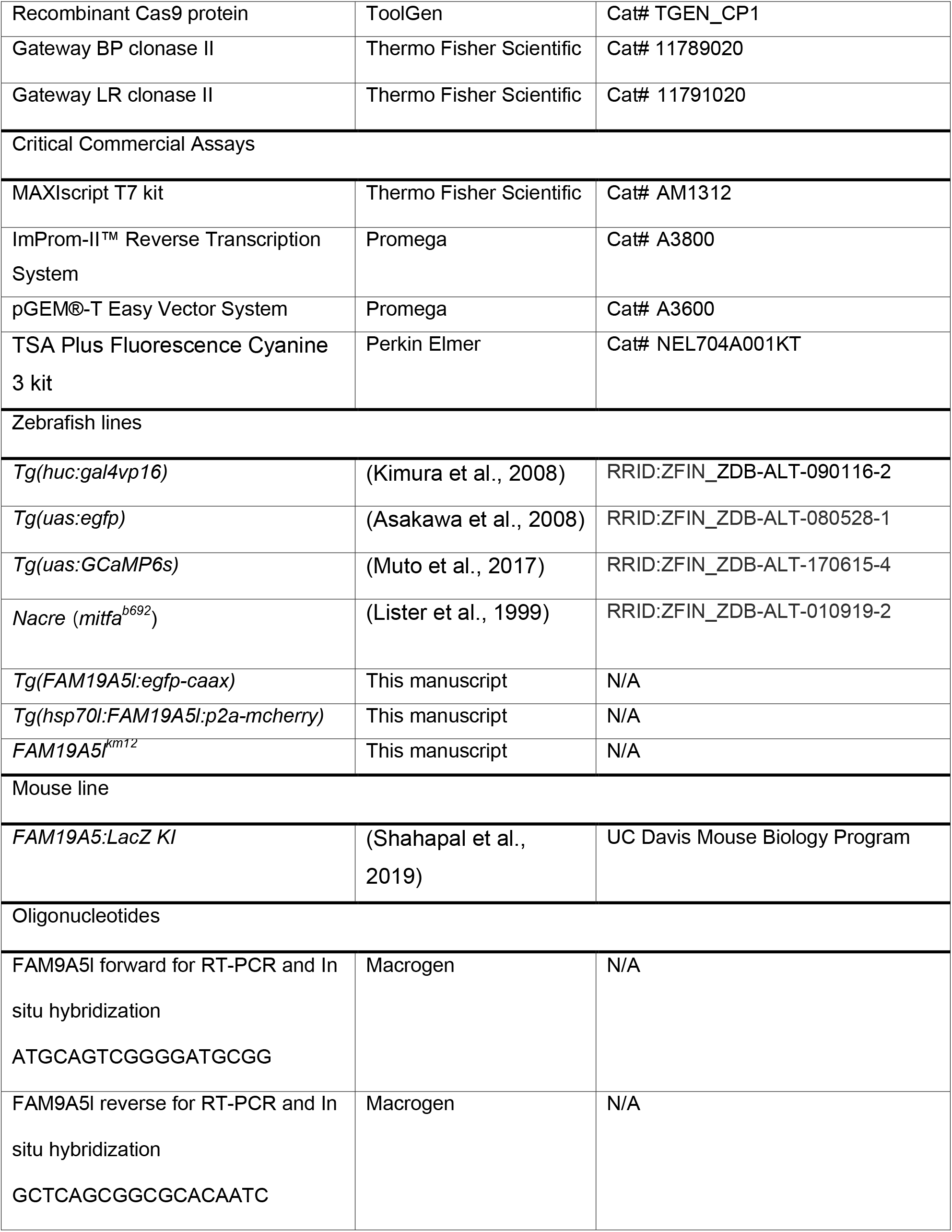
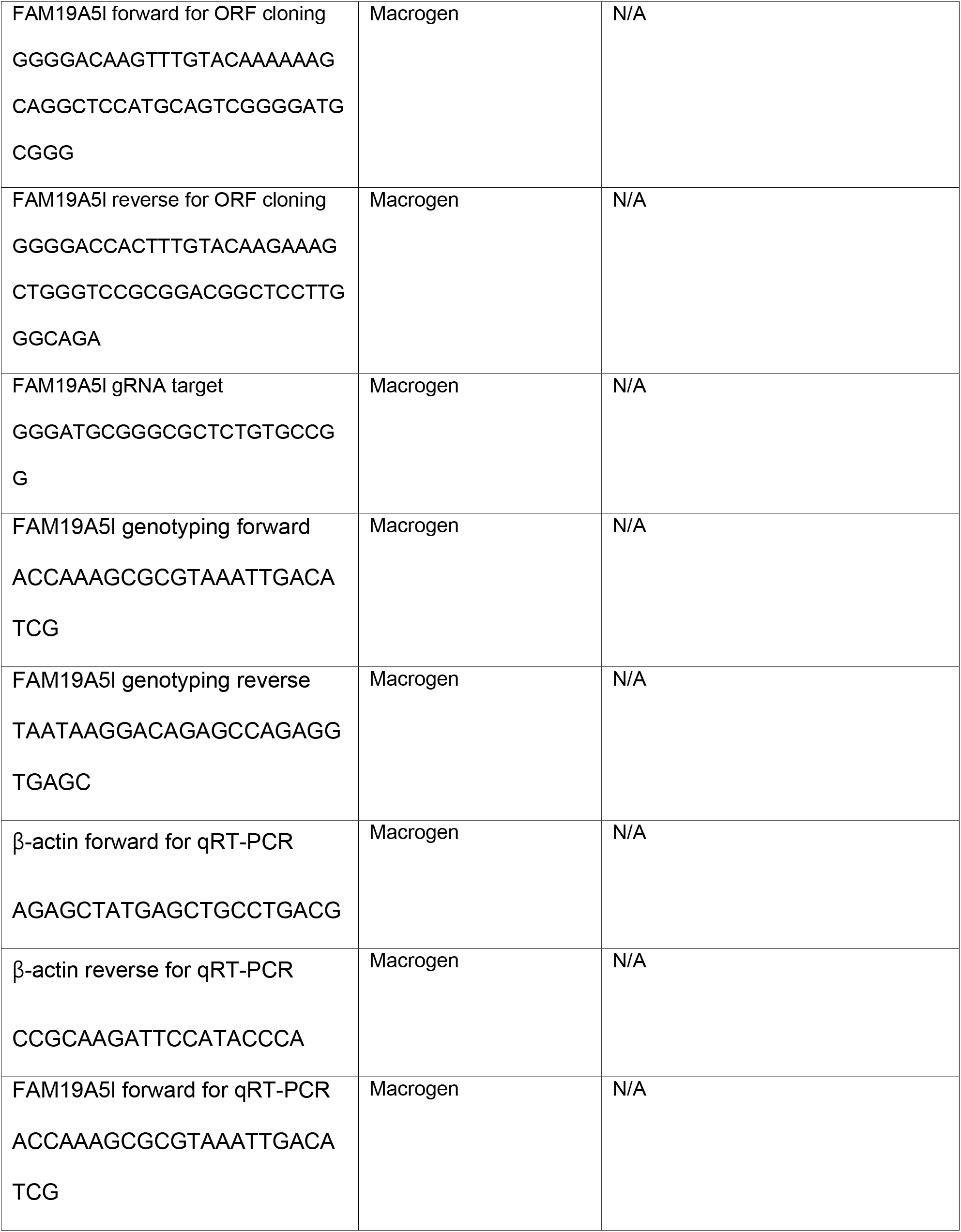
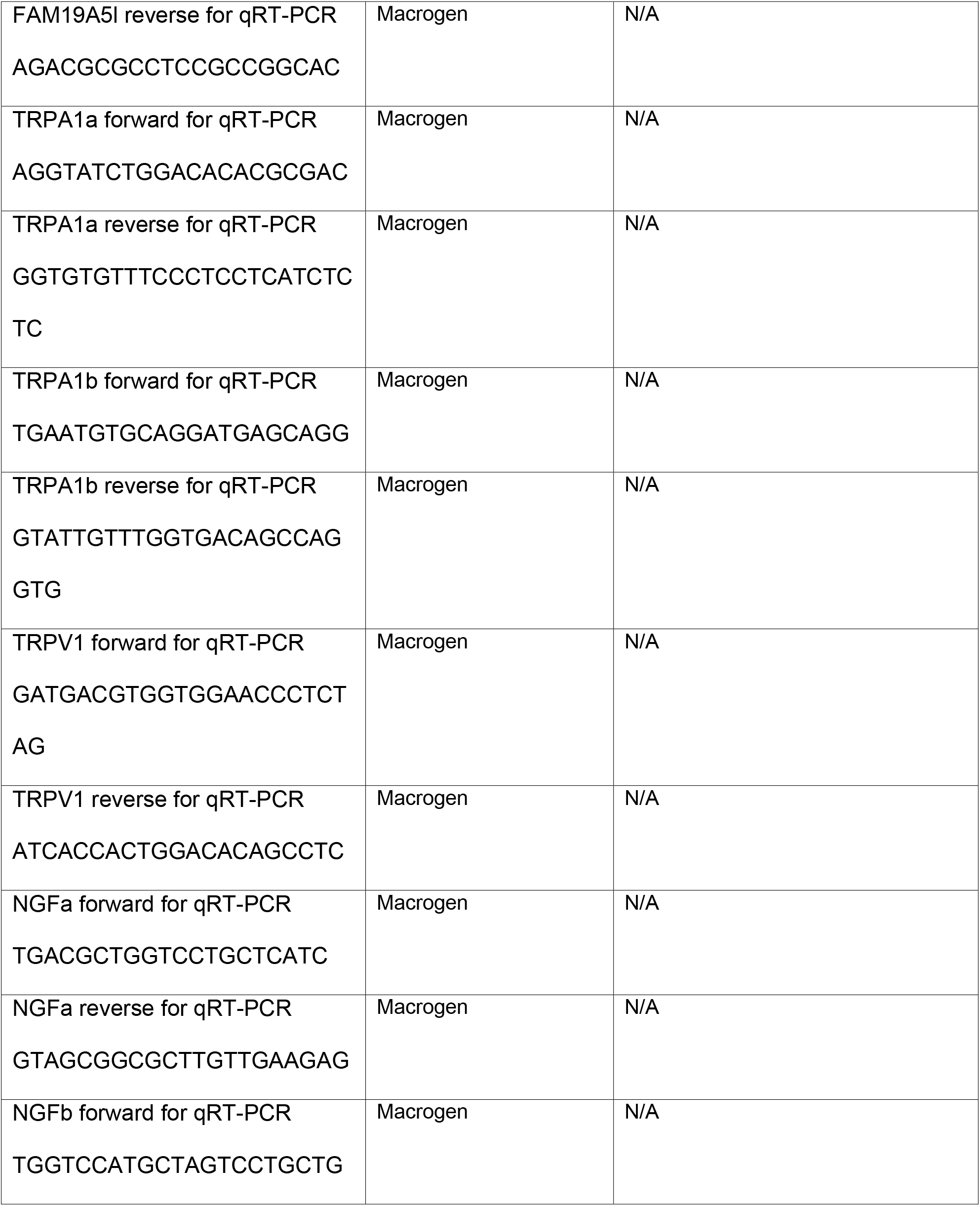
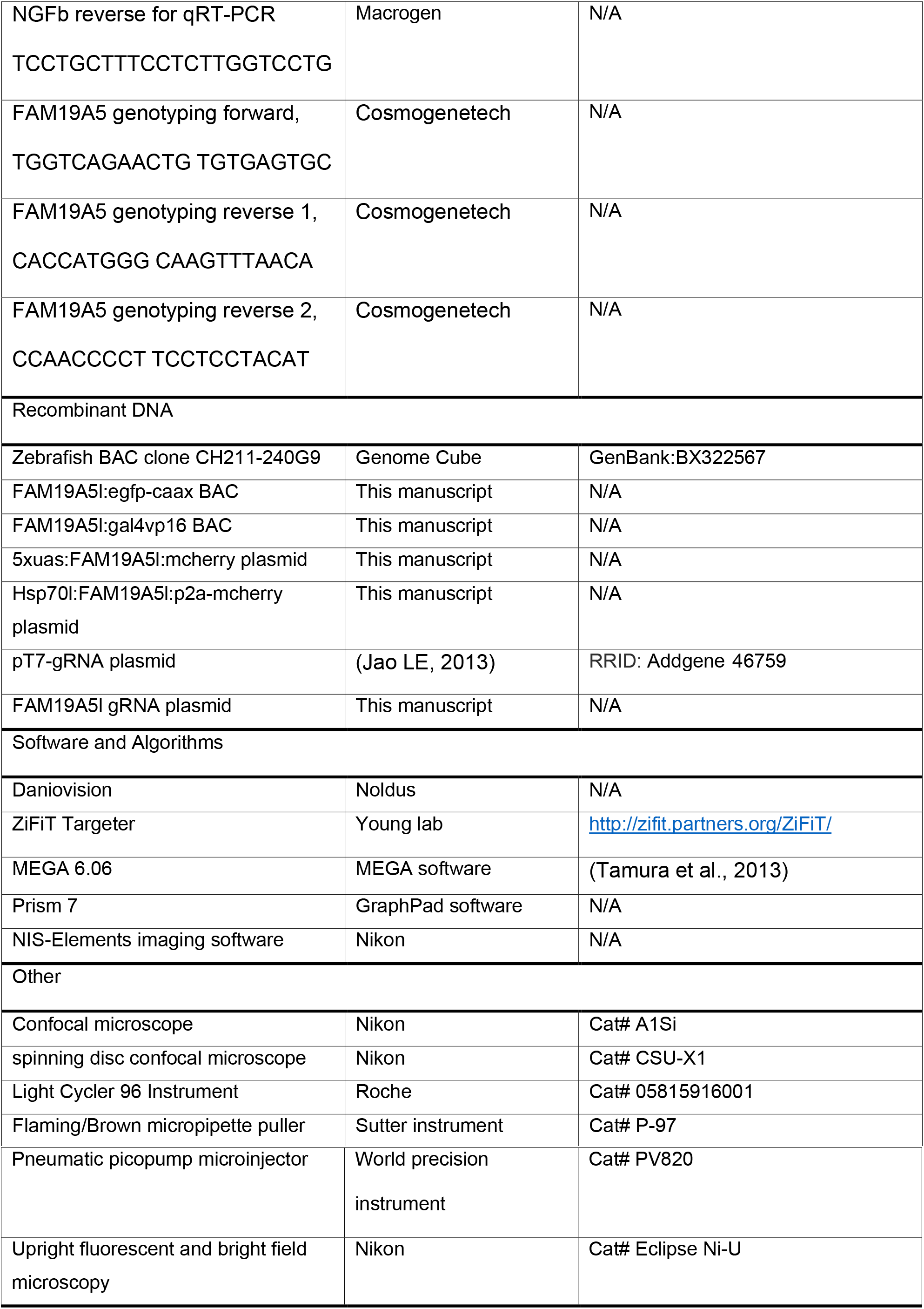

